# A single septin from a polyextremotolerant yeast recapitulates many canonical functions of septin hetero-oligomers

**DOI:** 10.1101/2024.05.23.595546

**Authors:** Grace E Hamilton, Katherine N Wadkovsky, Amy S Gladfelter

## Abstract

Morphological complexity and plasticity are hallmarks of polyextremotolerant fungi. Septins are conserved cytoskeletal proteins and key contributors to cell polarity and morphogenesis. They sense membrane curvature, coordinate cell division, and influence diffusion at the plasma membrane. Four septins homologs are conserved from yeasts to humans, the two systems in which septins have been studied most extensively. But there is also a fifth family of septin proteins that remain biochemically mysterious. Members of this family, known as Group 5 septins, appear in the genomes of filamentous fungi, and thus have been understudied due to their absence from ascomycete yeasts. *Knufia petricola* is an emerging model polyextremotolerant black fungus that can serve as a model system for understudied Group 5 septins. We have recombinantly expressed and biochemically characterized *Kp*AspE, a Group 5 septin from *K. petricola*, demonstrating that this septin––by itself *in vitro*–– recapitulates many of the functions of canonical septin hetero-octamers. *Kp*AspE is an active GTPase that forms diverse homo-oligomers, senses membrane curvature, and interacts with the terminal subunit of canonical septin hetero-octamers. These findings raise the possibility that Group 5 septins govern the higher order structures formed by canonical septins, which in *K. petricola* cells form extended filaments. These findings provide insight into how septin hetero-oligomers evolved from ancient homomers and raise the possibility that Group 5 septins govern the higher order structures formed by canonical septins.

**Significance Statement:**

- Septins are understudied cytoskeletal proteins. Here, we biochemically characterized *Kp*AspE, a novel Group 5 septin from a polyextremotolerant black fungus.
- *Kp*AspE in isolation recapitulates many properties of canonical septin hetero-octamers *in vitro* and interacts with the Cdc11, the terminal subunit of those octamers.
- These findings provide insight into how ancient septins may have evolved and diversified from homopolymers and suggest that many of the septin functions were present in the ancestral protein.

## Introduction

The dynamic protein filaments of the cytoskeleton allow cells to sense and change their shapes, an ability fundamental to their survival. Septins are perhaps the least-understood polymers of the cytoskeleton. But they participate in many essential cellular processes, including cytokinesis, membrane organization, polarization, and morphogenesis (Field et al., 1996; Gladfelter et al., 2005; Longtine et al., 2000; McMurray & Thorner, 2009). Unsurprisingly, septin dysfunction is linked with numerous human diseases, including male infertility and cancer (Dolat et al., 2014). Septins are conserved across eukaryotes (with the exception of land plants) (Nishihama et al., 2011; Shuman & Momany, 2021), but they are most intensively studied in the system in which they were discovered: *S. cerevisiae* (Hartwell, 1971).

In budding yeast, the five mitotic septins assemble stepwise into palindromic hetero-octamers, the terminal subunit of which may be either Cdc11 or Shs1 (Khan et al., 2018; Weems & McMurray, 2017) (Figure 1). Octamers contain two kinds of interfaces: NC-interfaces made up of the N- and C-termini of the two interacting septins; and G-interfaces that occur through the GTPase domains of both septins (Kim et al., 2012). All septins contain a central P-loop GTPase domain, although not all hydrolyze GTP (Versele & Thorner, 2004). GTP binding and hydrolysis rates affect oligomer composition (Weems et al., 2014), but GTP hydrolysis by septins has been difficult to directly detect in cells (Vrabioiu et al., 2004). Septin octamers are the predominant species in the cytoplasm of fungal cells (Bridges et al., 2014), but octamers at the plasma membrane self-assemble into diverse higher-order structures including filaments, bundles, bars, meshes, and collars (Bridges et al., 2014; DeMay et al., 2011; Hernández-Rodríguez et al., 2012; Hernández-Rodríguez et al., 2014). *In vivo* and *in vitro*, septins localize to regions of positive membrane curvature, using an amphipathic helix in the C-terminus of Cdc12 (Bridges et al., 2016; Cannon et al., 2019; Shi et al., 2023). The ability to sense membrane curvature has also been demonstrated in human septins (Bridges et al., 2016; Nakazawa et al., 2023).

**Figure 1.**
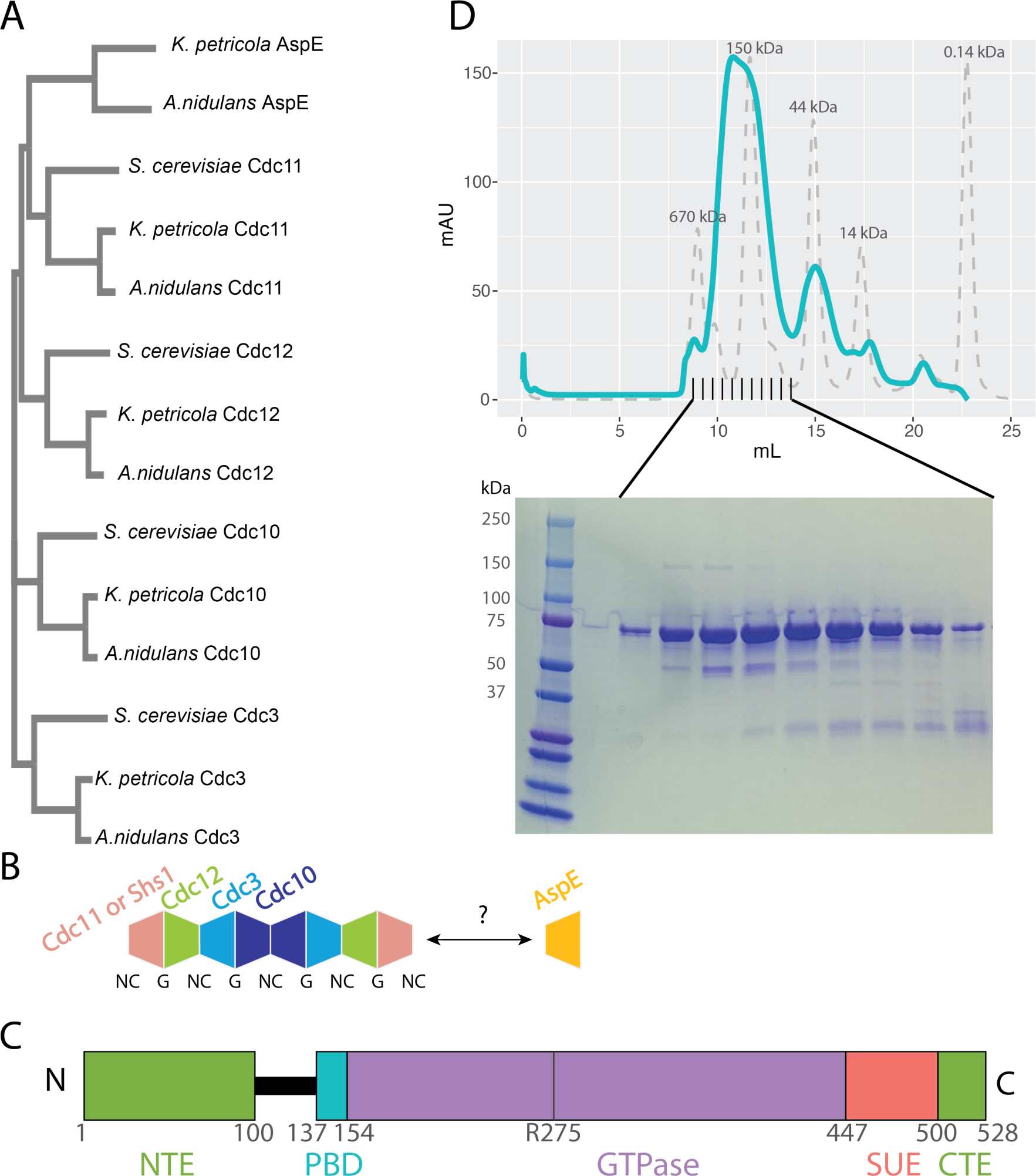
Identification and purification of a Group 5 septin from the emerging model fungus *K. petricola*. (A) Clustal Omega phylogenetic tree of representative fungal septins from *S. cerevisiae*, *A. nidulans*, and *K. petricola*. *S. cerevisiae* lacks an AspE homolog. (B) The palindromic structure of canonical septin hetero-octamers is made up alternating G- and NC-interfaces between septin subunits. (C) Predicted structural features of *Kp*AspE. Conserved features of all septins, such as the N-terminal extension (NTE), polybasic domain (PBD), GTPase domain, septin unique element (SUE), and C-terminal extension (CTE) are indicated with boxes. A vertical line indicates an arginine finger that is conserved in many non-opisthokont septins and some Group 5 septins. (D) Chromatogram of recombinantly expressed *Kp*AspE run on a Superdex 200 size exclusion chromatography (SEC) column and monitored at 260 nm as the final step of purification. Molecular weight standards analyzed on the same column are shown in gray. Below is a Coomassie-stained SDS-PAGE gel of the elution fractions. AspE has a predicted molecular weight of 73 kDa prior to cleavage of a C-terminal thioredoxin solubility tag, and the recombinant protein elutes from the Superdex column at a volume consistent with a dimer.

The number of septin genes varies between organisms, from as few as one in the green alga *Chlamydomonas reinhardtii* to as many as a dozen in humans (Shuman & Momany, 2021). Opisthokont septins are presently classified into five Groups, with members of the same group able to occupy orthologous positions in septin heteropolymers (Delic et al., 2024; Kinoshita, 2003; Shuman & Momany, 2021). For example, *S. cerevisiae* Cdc11 and Shs1 are both Group 3 septins, as one or the other occupies the terminal positions in every septin octamer. The extant diversity of septin orthologs and paralogs is thought to be due to a history of gene duplication and divergence (Auxier et al., 2019; Cannon et al., 2023). This suggests that septin octamers, like many hetero-oligomeric proteins, evolved from an ancestral homo-oligomer (Mallik & Tawfik, 2020). Thus, any modern septins that exist primarily as homo-oligomers may offer us a window into the evolutionary history of septins.

The vast majority of studies on opisthokont septins focus on members of Groups 1-4, which assemble into canonical hetero-octamers. Far less is known about Group 5 septins, which are found in filamentous fungi (Delic et al., 2024; Shuman & Momany, 2021). Septins have also been identified in the genomes of some non-opisthokonts, including ciliates, diatoms, chlorophyte algae and brown algae (Delic et al., 2024; Shuman & Momany, 2021; Vargas-Muñiz et al., 2016; Yamazaki et al., 2013). In the ciliate *Tetrahymena thermophila*, all three septins in its genome are Group 8 septins; they localize to and stabilize mitochondria (Wloga et al., 2008). The green alga *Chlamydomonas reinhardtii* has only one septin gene, the product of which is a Group 6 septin known as *Cr*SEPT. Crystal structures of *Cr*SEPT reveal that it dimerizes across a G-interface and that a catalytic arginine residue is essential for dimerization (Pinto et al., 2017). Intriguingly, this arginine finger, which is present in the majority of non-opisthokont septins, is also present in some Group 5 septins, and they are the only group of opisthokont septins with this feature (Delic et al., 2024). To our knowledge, there is no structural data on Group 5 septins, not even in the model filamentous fungus *Aspergillus nidulans*, in which the Group 5 septin known as AspE was first described (Hernández-Rodríguez et al., 2014; Momany et al., 2001). *In vivo* studies of *An*AspE suggest that it does not incorporate into heteropolymers with the canonical septins of Groups 1-4, although it was required for their assembly into higher-order structures during multicellular development (Hernández-Rodríguez et al., 2014). Biochemical data on fungal Group 5 septins is lacking, despite the importance of septins in the virulence of plant and human fungal pathogens (Boyce et al., 2005; Chen et al., 2016; Dagdas et al., 2012; Kozubowski & Heitman, 2010; Warenda et al., 2003).

In order to address this gap in our knowledge, we turned to black yeasts, a group of melanized ascomycetes that have adapted to survive in diverse extreme environments and exhibit dramatic plasticity in their morphology and mode of cell division (Goshima, 2022; Mitchison-Field et al., 2019; Nai et al., 2013; Ruibal et al., 2009; Réblová et al., 2013). *Knufia petricola* is an emerging model black fungus for which genetic tools have been developed (Erdmann et al., 2022; Voigt et al., 2020). In addition to orthologs of the canonical septins (members of Groups 1-4), we identified a Group 5 septin in the genome of *K. petricola*. We recombinantly expressed and purified this septin, dubbed *Kp*AspE (for its *Aspergillus* ortholog). *in vitro*, *Kp*AspE recapitulates many of the features of canonical septin hetero-oligomers. Gel filtration and mass photometry indicate that *Kp*AspE forms homo-oligomers in a pH-dependent manner. In a membrane curvature-sensing assay, *Kp*AspE appears able to both bind supported lipid bilayers and discriminate between different degrees of positive membrane curvature– in a similar manner to heteropolymeric septins. Of the four canonical *Knufia* septins, only *Kp*Cdc11 was able to interact with Kp*AspE* in a yeast two-hybrid assay. Taken together, these results suggest that KpAspE homo-oligomers may replicate some of the most vital functions of canonical septin hetero-oligomers. Additionally, we found that *K. petricola* undergoes a dimorphic switch to pseudohyphal growth in response to carbon starvation and that the canonical septin *Kp*Cdc12 localizes to thick filaments in both round and pseudohyphal cells.

## Results

### Identification and purification of a *K. petricola* Group 5 septin

In order to begin characterization of the septin cytoskeleton in the emerging model fungus *K. petricola*, we used the basic local alignment search tool (BLAST)(Altschul et al., 1990) to search the published genome (GenBank assembly GCA_002319055.1) for homologs of the canonical budding yeast septins. We identified five septin genes in the *K. petricola* genome, four of which were orthologous to the four canonical mitotic septins of budding yeast (Figure 1A) These septins (Cdc11, Cdc12, Cdc3, Cdc10) assemble to palindromic hetero-octamers in *S. cerevisiae* (Khan et al., 2018; Weems & McMurray, 2017) (Figure 1B). We also identified a fifth *K. petricola* septin that resembled the Group 5 septin AspE from model filamentous fungus *Aspergillus nidulans* more closely than it did any budding yeast septin (Figure 1A). The predicted *K, petricola* protein shared 55.93% sequence identity with *An*AspE. Of the *S. cerevisiae* septins, *Sc*Cdc11 showed the greatest sequence identity with *Kp*AspE (22.74%).

All septins share a conserved structure: a central GTPase domain flanked by a polybasic domain (PBD) and the septin unique element (SUE). This conserved core is in turn flanked by N- and C-terminal extensions whose lengths vary dramatically between septins (Shuman & Momany, 2021). Thus, the start- and endpoints of the *K. petricola* septins genes could not be deduced from simple alignment with orthologous gene sequences. We used the AUGUSTUS software to predict initiation sites, stop codons, and splice sites for all five *K. petricola* septin genes (Stanke & Morgenstern, 2005). In order to confirm these predictions experimentally, we used rapid amplification of cDNA ends (RACE) (Frohman, 1993) (Supp. Table 1). Like most Group 5 septins, *Kp*AspE contains a relatively long NTE (Figure 1C). Every *K. petricola* septin gene contained at least one intron (Supp. Table 2), and the positions of some introns were conserved relative to the orthologous *A. nidulans* genes (Supp. Figure 1). Exon locations were corroborated by comparison to an independently collected RNA-seq dataset (Julia Schumacher, personal communication.)

In order to biochemically characterize the *Kp*AspE protein, we codon-optimized the coding sequence of the gene for *E. coli* and incorporated this sequence into an expression vector with a cleavable N-terminal 6xHis-thioredoxin tag, for affinity purification and enhanced yield and solubility, respectively (Savitsky et al., 2010). The recombinantly expressed protein was predicted to have a molecular weight of 72.97 kDa prior to TEV cleavage of the N-terminal tag and a molecular weight of 59.04 kDa after cleavage. *Kp*AspE was purified by nickel affinity chromatography followed by size exclusion chromatography, where it eluted from the column at about the same volume as a 150-kDa protein standard, consistent with the un-cleaved protein behaving primarily as a dimer in solution (Figure 1D).

We next examined the biochemical and biophysical properties of *Kp*AspE protein to evaluate its behavior compared to canonical septin hetero-octamers.

### *Kp*AspE is an active, slow GTPase

Septins bind and in some cases hydrolyze GTP. Of the canonical *S. cerevisiae* septins, Cdc10 and Cdc12 are active GTPases *in vitro*, while Cdc3 and Cdc11 are not (Versele & Thorner, 2004). Based on the presence of conserved residues involved in nucleotide binding (D329) and hydrolysis (R275), we hypothesized that *Kp*AspE has GTPase activity.

In order to test for GTPase activity of *Kp*AspE, we first asked if this protein could bind and exchange guanosine nucleotides. We used a fluorescent analog of GDP, BODIPY-GDP, which emits light only when protein binding eliminates quenching between the adenine and BODIPY moieties of the molecule (Figure 2A). The addition of *Kp*AspE caused a rapid increase in fluorescence, followed by a slow decrease, consistent with the rapid binding and exchange of BODIPY-GDP by the protein (Figure 2B.)

**Figure 2.**
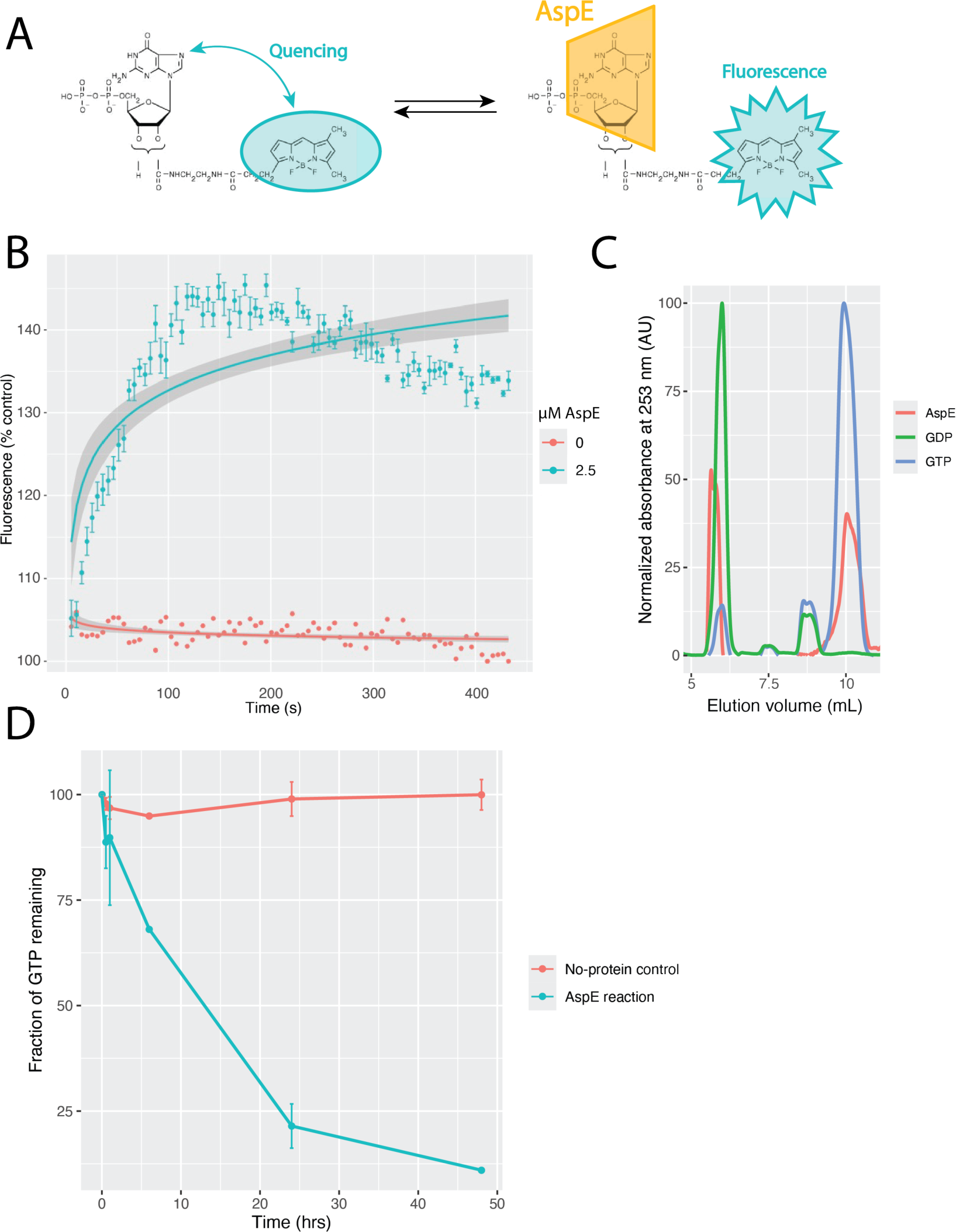
AspE binds and hydrolyzes GTP. (A) Schematic of the BODIPY-guanine nucleoside binding assay. (B) BODIPY-GDP binding and exchange. The results of three technical replicates are shown, with error bars indicating standard error. (C) Elution profile of *Kp*AspE’s co-purifying nucleotides and nucleotide standards from the Hypersil™ ODS C18 HPLC column. Recombinant *Kp*AspE protein contains a mixture of bound GDP and GTP. (D) The conversion of GTP into GDP by KpAspE was monitored at room temperature by HPLC. The results of two technical replicates are shown; error bars indicate standard error at each timepoint.

Curious about the nucleotide-binding state of our recombinantly expressed and purified *Kp*AspE, we boiled samples to denature the protein and separate it from co-purifying nucleotides, then analyzed those samples using high-performance liquid chromatography (HPLC). We observed two elution peaks, which coincide with GDP and GTP standards (Figure 2C). We conclude that recombinant *Kp*AspE protein contains a mixture of bound GDP and GTP.

We next asked if purified KpAspE could hydrolyze GTP *in vitro*. GTPase activity assays were performed by incubating *Kp*AspE with a ten-fold excess of GTP for different times and evaluating the GTP and GDP content of the sample by HPLC. Over 48 hours, nearly all of the GTP was hydrolyzed in the sample containing *Kp*AspE, but not in a control reaction lacking *Kp*AspE (Figure 2D). We conclude that *Kp*AspE is an active GTPase, albeit a very slow one under the *in vitro* conditions of this assay. Based on the rapid nucleotide exchange observed with BODIPY-GDP, we further conclude that the hydrolysis reaction itself–– rather than nucleotide exchange–– is the rate-limiting factor in *Kp*AspE’s slow hydrolysis of GTP.

### *Kp*AspE forms homo-oligomers through distinct NC- and G-interfaces

Based on the size exclusion chromatography performed as the final step of protein purification, *Kp*AspE behaves primarily as a dimer at micromolar concentrations in purification buffer (300 mM KCl). The concentration of soluble septins in the cytoplasm has been previously measured at 100-200 nM for three different fungal species: *S. cerevisiae*, *S*. *pombe*, and *A. gossypii* (Bridges et al., 2014). In *A. nidulans*, *An*AspE is reported to be less abundant than Groups 1-4 septins (Hernández-Rodríguez et al., 2014). In order to determine the oligomeric state of purified *Kp*AspE protein in a nanomolar concentration regime and under varying buffer conditions, we turned to mass photometry.

The predicted molecular weight of KpAspE, after cleavage of the N-terminal 6xHis-thioredoxin tag, is 59.045 kDa. Analysis of *Kp*AspE by mass photometry at pH 7.4, at low nanomolar concentrations, yielded two majors peaks: one at about 60 kDa and the other at 120 kDa (Figure 3). This suggest that under these conditions, *Kp*AspE is found as both a monomer and a dimer, in about equal abundance (Figure3). The presence of a very small peak between 360 and 420 kDa may indicate the presence of larger oligomers, albeit in very low abundance.

**Figure 3.**
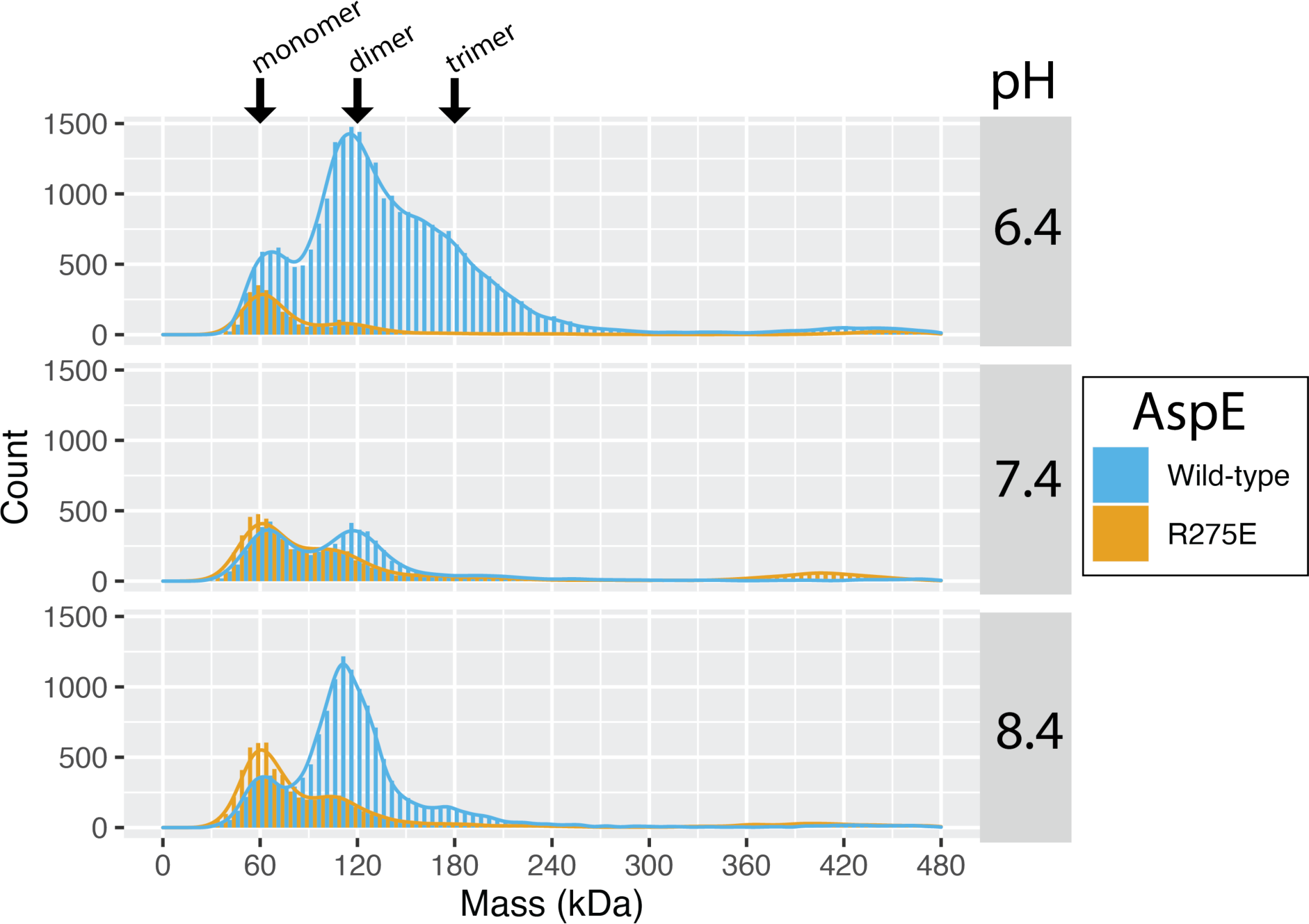
AspE form diverse homo-oligomers in solution. Mass photometry at pH 6.4, 7.4, and 8.4. The predicted molecular weight of *Kp*AspE, after cleavage of the N-terminal 6xHisthioredoxin tag, is 59.045 kDa.

We hypothesized that *Kp*AspE was dimerizing across a G-interface made up primarily of its GTPase domain, as has been previously reported for an algal Group 6 septin (Pinto et al., 2017). Pinto and colleagues report a charge reversal mutation to the catalytic arginine finger at the G-interface of *Chlamydomonas reinhardtii*’s Group 6 septin completely abrogates dimer formation (Pinto et al., 2017). This arginine residue is conserved in some Group 5 septins, including *Kp*AspE (Delic et al., 2024), so we made the analogous mutation: R275E. This mutation did not completely eliminate dimer formation by *Kp*AspE, but it did shift the distribution of the population to monomers as the more abundant species (Figure 3). This suggests some contribution of the catalytic arginine in stabilizing the G-interface for dimerization.

Previous work has shown that filament formation by canonical septin octamers is sensitive to solution pH (Jiao et al., 2020). In order to determine if *Kp*AspE-*Kp*AspE interactions are similarly sensitive, we varied the pH in our mass photometry experiments. At pH 6.4, we observed a third species with a molecular weight of approximately 180 kDa, consistent with the formation of a *Kp*AspE trimer in solution, in addition to the previously observed dimer (∼120 kDa) and monomer (∼60 kDa) peaks (Figure 3). Because dimer formation appeared to occur across a G-interface, trimer formation must occur across either a NC-interface or some other interface that has not been previously described for septins. However, notably, the R275E mutation still abrogated trimer formation at pH 6.4, causing the monomer to be the most prevalent species. This suggests that dimer formation via the G-interface may be a requirement for trimer formation. We then assessed the effects of increased pH and found that at pH 8.4, a dimer peak predominated, although smaller monomer and trimer peaks were also present for WT *Kp*AspE. The R275E mutation eliminated trimers, and it rendered monomers more abundant than dimers at pH 8.4. Thus, we see that the equilibria of AspE protein between monomers, dimers and trimers is sensitive to pH and may involve an ordered assembly beginning with a G-interface mediated dimer.

Recombinantly expressed and purified canonical septin octamers form filaments on supported lipid bilayers *in vitro* (Bridges et al., 2014; Khan et al., 2018; Szuba et al., 2021). Under the same conditions that promote filamentation of *S. cerevisae* and human canonical septins on synthetic bilayers (5-100 nM protein; pH 7.4; 25-75 mM KCl), we did not observe any filamentation of fluorescently labeled *Kp*AspE (Supp. Figure 2). This could indicate that polymerization into long filaments is not a function of this septin, or that conditions, cofactors, and/or binding partners necessary for filamentation of *Kp*AspE were not present in this assay.

### *Kp*AspE exhibits a membrane curvature preference that is shallower than that of canonical septins

We next asked if *Kp*AspE has the ability to sense micron-scale positive membrane curvature, a feature of canonical septin oligomers. Presented with beads of different sizes coated in supported lipid bilayers, both mammalian and budding yeast canonical septins preferentially adsorb to 1-µm diameter beads reflecting a micron-scale curvature preference (Bridges et al., 2016). Filament formation is not required for this curvature preference in some contexts (Bridges et al., 2016), so we subjected purified *Kp*AspE to the same curvature preference assay (Figure 4A-B). Presented in the same reaction with three beads of different sizes (0.3, 1, and 5 µm in diameter), *Kp*AspE preferentially adsorbed to the shallowest membrane curvature present in the assay, i.e. to the largest beads (Figure 4). This is in contrast to *S. cerevisiae* hetero-oligomeric septin complexes in a similar assay, where the highest absorption would be seen on the 1 µm diameter beads (Shi et al., 2023). Because *Kp*AspE preferentially adsorbed to the largest beads present in the assay, it is possible that *Kp*AspE actually prefers membrane curvature that is shallower than any present in this experiment.

**Figure 4.**
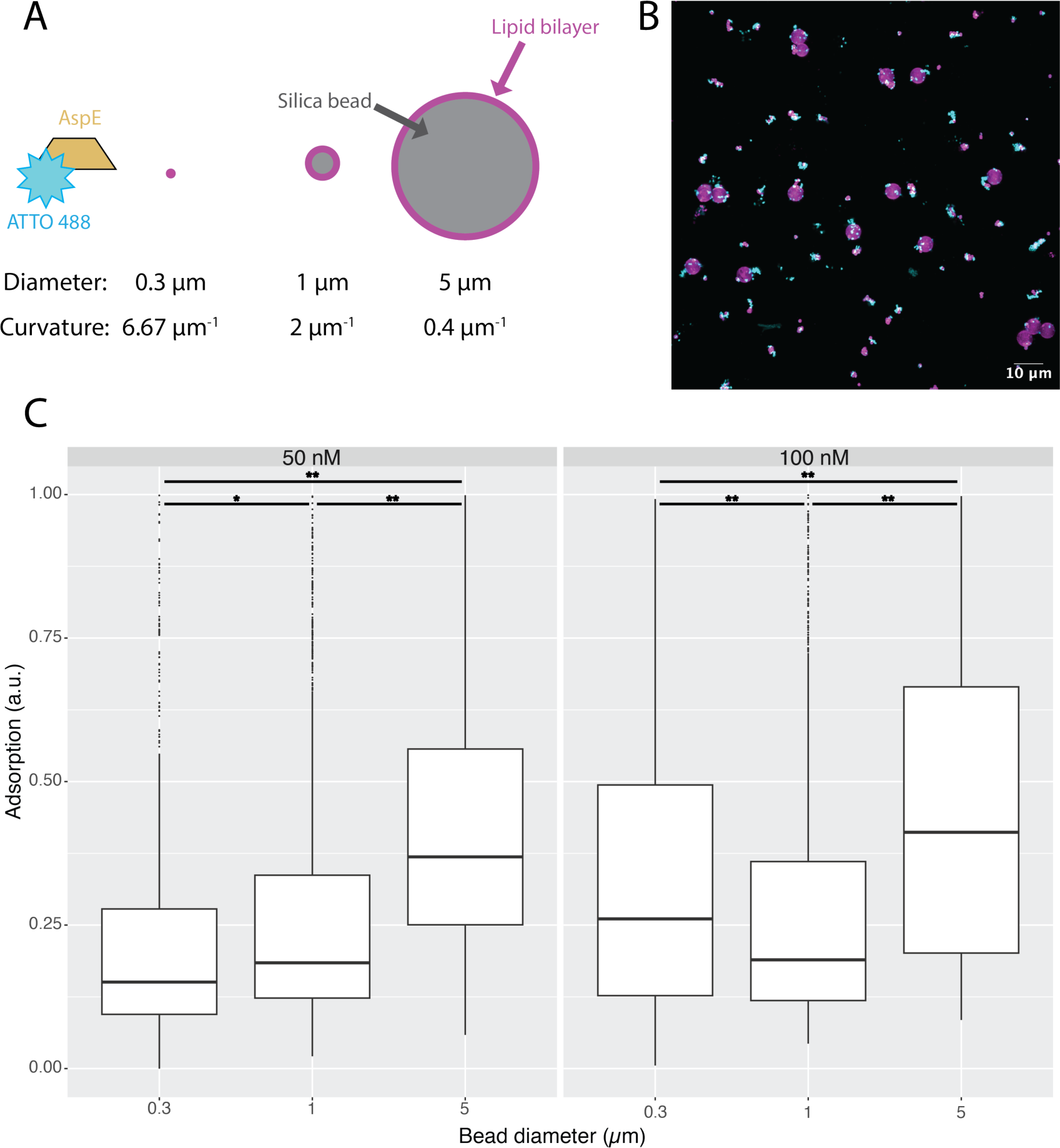
*in vitro* AspE preferentially localizes to shallow positively curved membranes. (A) Schematic of assay. Supported lipid bilayer (25% PI and 75% PC) and trace Rh-PE–coated silica beads ranging from 0.3 to 5 μm in diameter mixed with either 50 nM or 100 nM *Kp*AspE labeled with Atto 488 NHS ester for visualization. (B) Microscopy images of beads. The Rh-PE is shown in magenta, and *Kp*AspE-Atto488 is shown in green. (C) Boxplot of AspE adsorption levels observed for beads of different sizes. Boxes extend from the lower quartile to the upper quartile. Whiskers extend to 1.5 times the interquartile range beyond each quartile. A Kruskal-Wallis test with pairwise Wilcox post-hoc tests was used to compare distributions and calculate p-values. ** indicates p < 0.005; * indicates p < 0.05. A plot of individual datapoints, colored by biological and technical replicate can be found in Supplementary Figure 3.

### *Kp*AspE interacts with only the Cdc11 canonical septin

We next asked whether or not Kp*AspE* could directly interact with any of the canonical septins also present in *K. petricola*. The lack of a septin antibody that cross-reacts with *K. petricola* septins prompted us to turn to a yeast two-hybrid assay to assess interactions. In a yeast two-hybrid assay, *Kp*AspE showed interaction with only one other *Knufia* septin: *Kp*Cdc11 (Figure 5). This interaction could be abrogated by either deleting the C-terminal extension (amino acids 299-380) of *Kp*Cdc11 or by deleting the N- or C-terminal extensions of *Kp*AspE (amino acids 1-100 and 502-528, respectively) (Figure 5B). These results were consistent regardless of which septin was fused to the activating domain and which was fused to the DNA binding domain of the Gal4 transcription factor (Supp. Figure 3). Removing the N- or C-terminus of *Kp*AspE did not prevent it from interacting with itself in the yeast two-hybrid assay (Supp. Figure 3), consistent with our hypothesis that *Kp*AspE dimerizes across a G-interface but interacts with *Kp*Cdc11 across a NC-interface. In order to assess the plausibility of this hypothesis, AlphaFold Multimer was used to predict the structure of a *Kp*AspE homodimer (Evans et al., 2022). Consistent the data from our yeast two-hybrid assays, AlphaFold predicted that *Kp*AspE dimerizes across a G-interface (Figure 5D). We next used AlphaFold to predict the structure of a *Kp*AspE-KpCdc11 heterodimer. AlphaFold also predicted that these two septins would dimerize across a G-interface (Figure 5E). This seems unlikely given our finding that the N- and C-termini of *Kp*AspE are necessary for its interaction with *Kp*Cdc11. Taken together, these results raise the possibility that in some circumstances *Kp*AspE may engage in hetero-oligomers.

**Figure 5.**
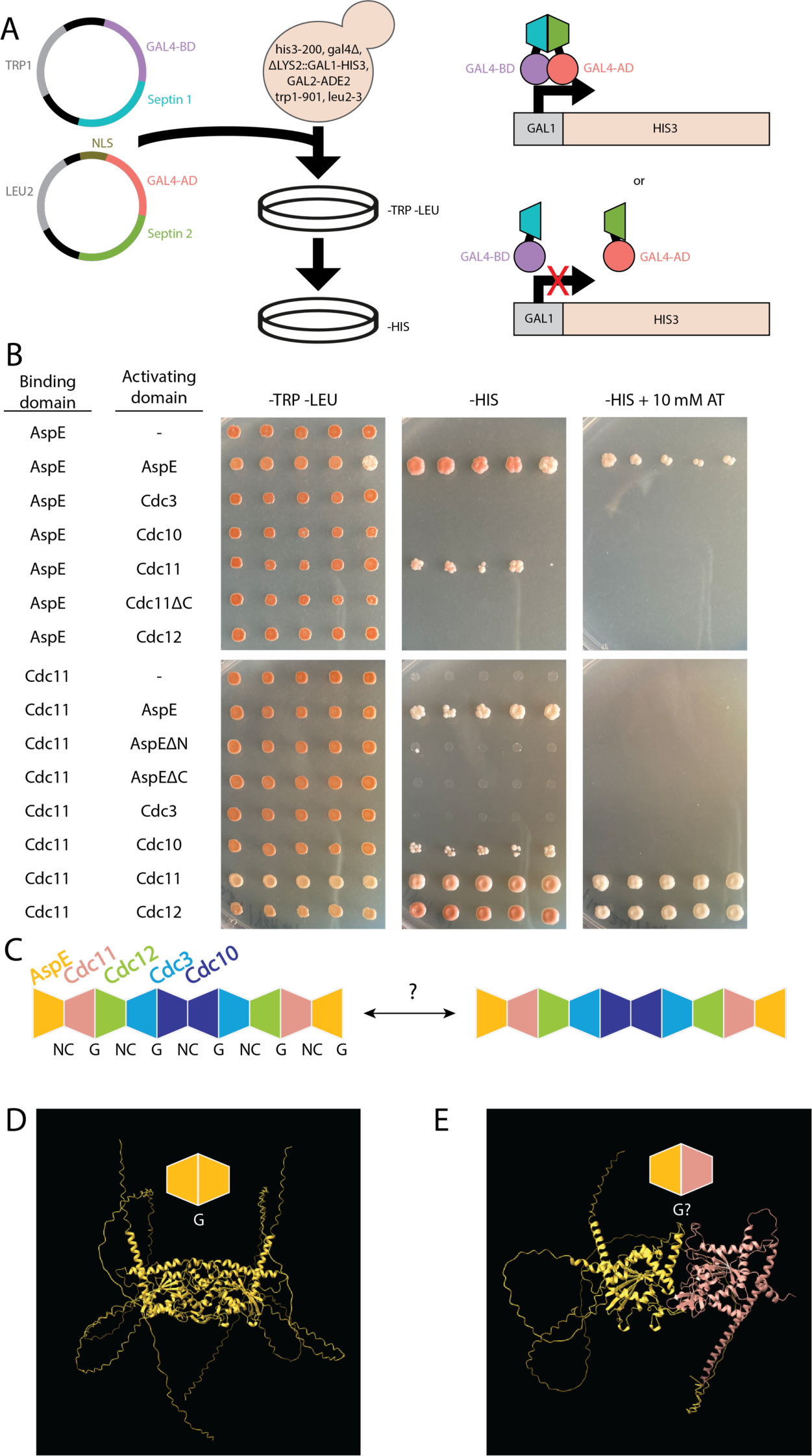
Of the Groups 1-4 septins, only Cdc11 is capable of binding AspE. (A) Schematic of yeast two-hybrid assay. One candidate protein was fused to the C-terminus of the activating domain (AD) of the transcription factor Gal4, while the other candidate protein was fused to the C-terminus of Gal4’s DNA-binding domain (BD). Both the HIS3 and ADE2 genes are under the control of GAL promoters. Thus, both beige coloration and growth on media lacking histidine are indicative of physical interaction between the septin proteins fused to the DNA-binding and activating domains of the transcription factor Gal4. (B) Yeast two-hybrid assay results. (C) Schematic of conclusions drawn from yeast two-hybrid assays. In this *in vivo* assay, *Kp*AspE is capable of interacting with both itself and with *Kp*Cdc11. This suggests that *Kp*AspE could either cap septin octamers or mediate interactions between them. (D) AlphaFold-predicted structure of a *Kp*AspE-*Kp*AspE homodimer. (E) AlphaFold-predicted structure of a *Kp*AspE-*Kp*Cdc11 heterodimer.

### *K. petricola* transitions pseudohyphal growth in response to carbon starvation

We next asked if *K. petricola* exhibited cell shape plasticity, which is a hallmark of polyextremotolerant fungi and is likely related to septin function. Culturing wild-type *K. petricola* under a variety of conditions, we noted that colonies grown on solid synthetic complete media exhibited smooth edges, while those grown on water agar had fuzzy, ragged colony borders (Figure 6A). Hypothesizing that the lack of a specific nutrient might trigger this change in colony morphology, we then cultured wild-type *K. petricola* on solid synthetic complete media lacking either nitrogen, phosphorus, sulfur, or carbon. Only under carbon starvation conditions did *K. petricola* colonies recapitulate the fuzzy morphology seen on water agar (Figure 6A). Examining these cells under a microscope, we noted that *K. petricola* cells grown on rich or synthetic complete media are nearly spherical, while those grown under carbon starvation conditions form extended end-to-end chains of more elongated cells (Figure 6B). Based on the presence of calcofluor stain between the cells, indicative of septa, they do not share a continuous cytoplasm, making this pseudo-hyphal growth, rather than the formation of true hyphae.

**Figure 6.**
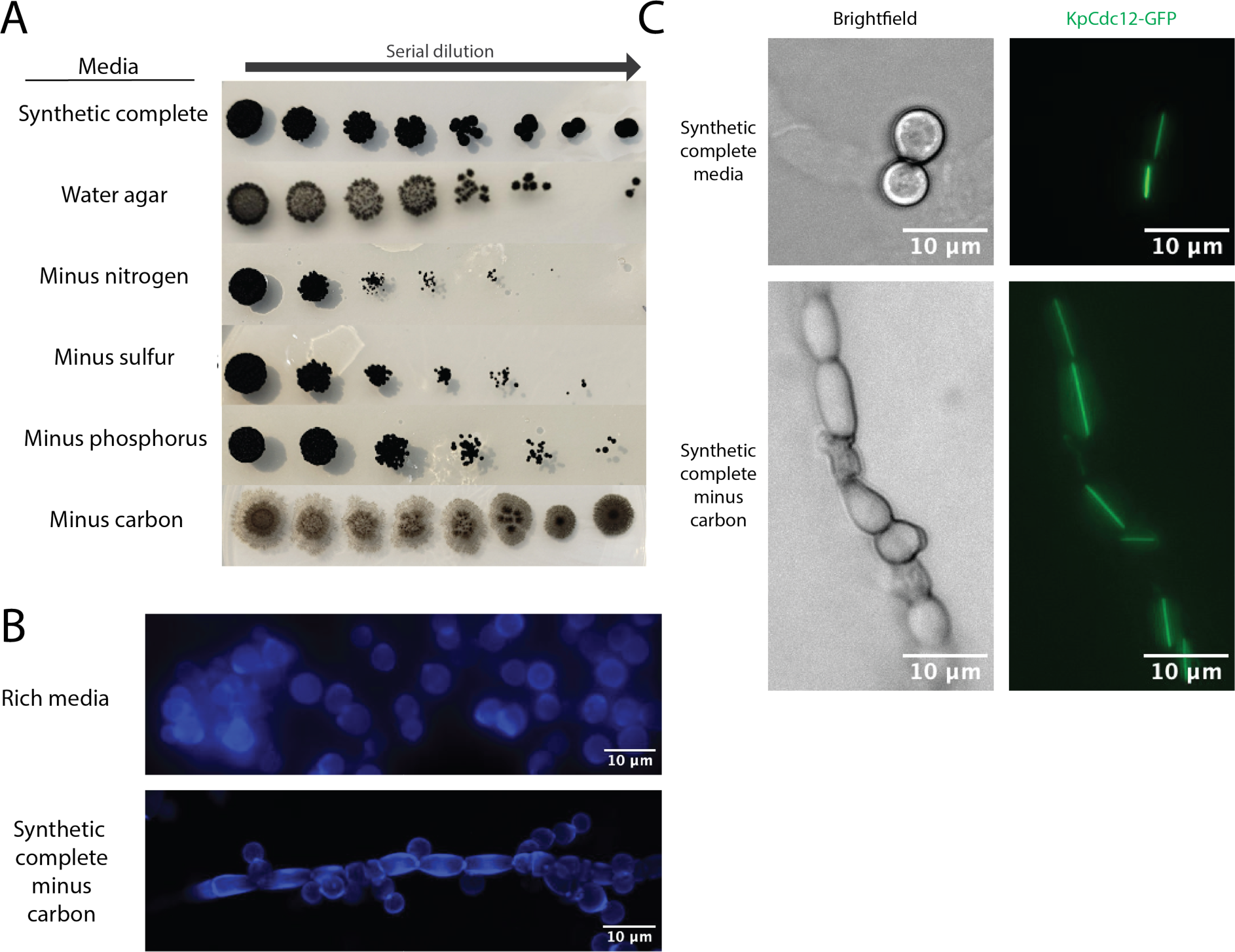
Septin localization and cell shape plasticity in response to carbon starvation in *K. petricola*. (A) Representative images of *K. petricola* colony morphology on synthetic complete and dropout media. (B) Fluorescence microscopy images of calcofluor white-stained *K. petricola* cells grown under standard or carbon-starved conditions. (C) Fluorescence microscopy images of *Kp*Cdc12-GFP signal in *K. petricola* colonies grown on synthetic complete or carbon dropout media.

### Cdc12 localizes to filaments that span the length of *K. petricola* cells

Given the established role of septin proteins in controlling cell shape, we next asked whether the localization patterns of *K. petricola* septins differed between round and pseudohyphal cells. In order to determine the localization patterns of canonical septins, we tagged the *K. petricola* Group 4 septin, *Kp*Cdc12, with an N-terminal GFP tag at its endogenous locus. We then used fluorescence microscopy to examine the localization pattern of this septin in cells grown under nutrient-rich and carbon starvation conditions. In both cases, *Kp*Cdc12 localizes exclusively to thick filaments that span the entire length of the cell (Figure 6C). It is unclear if this assembly is the native organization or potentially tag-induced, but the cells showed growth and shapes comparable to wild-type indicating that the tagged protein is likely functional. In the case of elongated, pseudohyphal cells, the *Kp*Cdc12 filaments align with the long axis of the cell. We also labeled *Kp*AspE with GFP at its endogenous locus, but no clear localization pattern was observed (data not shown), perhaps because this septin is expressed at lower levels than the canonical four septins (Hernández-Rodríguez et al., 2014).

## Discussion

We have recombinantly expressed and biochemically characterized AspE, a Group 5 septin from the emerging model black yeast *Knufia petricola*, demonstrating that this septin––by itself *in vitro*–– recapitulates many of the properties of canonical septin hetero-octamers. *Kp*AspE is an active GTPase capable of forming diverse homo-oligomers, sensing membrane curvature, and interacting with the terminal subunit of canonical septin hetero-octamers. To our knowledge, this is the first biochemical characterization of a Group 5 septin.

While septins are well-established as GTP-binding proteins, the functional role of GTP-binding and hydrolysis has been a challenge to decipher. Nucleotide binding state has been shown to regulate the assembly of septin homo-oligomers and hetero-oligomers (Weems & McMurray, 2017; Weems et al., 2014; Zent & Wittinghofer, 2014), however across a yeast cell cycle, GTP hydrolysis was not detectable, suggesting that there may not be further roles in higher-order assemblies (Vrabioiu et al., 2004). Taken together, our mass photometry and yeast two-hybrid assays support the idea that *Kp*AspE forms homodimers across its nucleotide-binding G-interface and interacts with *Kp*Cdc11 across a NC-interface (Figures 2 and 5). Our finding that *Kp*AspE is an active, slow GTPase therefore has profound implications for how this septin interacts with both itself and with canonical septins. Recombinantly expressed *Kp*AspE copurifies with a mixture of GTP and GDP, in contrast to the Group 6 septin *Cr*SEPT, which was found to be free of any bound nucleotide when expressed recombinantly in *E.coli* (Pinto et al., 2017). *Kp*AspE also hydrolyzes GTP to GDP at a rate much slower than that of *Cr*SEPT, despite conservation of the arginine finger motif (Pinto et al., 2017). In budding yeast, slow GTP hydrolysis by *Sc*Cdc12 allows stable subpopulations of GDP- and GTP-bound monomers, which are preferentially recruited to form hetero-oligomers with alternate binding partners (*Sc*Cdc11 or *Sc*Cdc3) (Weems & McMurray, 2017). It is tempting to speculate that *Kp*AspE interactions with itself and *Kp*Cdc11 might be regulated in a similar manner. Another possibility is that interactions at the NC-interface regulate nucleotide hydrolysis at the G-interface, as has been observed for *Sc*Cdc3, a budding yeast septin that–– like *Kp*AspE–– has a notably long N-terminal extension that is predicted to be largely disordered (Weems & McMurray, 2017). Thus, it seems likely that either the nucleotide-binding state of *Kp*AspE regulates its oligomeric state or the opposite. Further study will be needed to distinguish between these two possibilities.

Another factor that can govern the oligomeric state of *Kp*AspE is pH. Lowering the pH to 6.4 greatly increased the prevalence of trimers in mass photometry experiments (Figure 3), suggesting a protein-protein interface containing critical protonated histidine residues. We postulate that this pH-sensitive second interaction surface is a NC-interface. Indeed, we see several histidine residues at the predicted NC-interface: H122, H210, and H407. However, repeated attempts to purify a triple histidine-to-alanine mutant were unsuccessful, apparently due to the formation of large aggregates or homo-oligomers that eluted in the void volume during size exclusion chromatography (data not shown). This suggests that disruption of the NC-interface either destabilizes the protein or allows uncontrolled self-association. Moreover, the exceptionally long N-terminal extensions of Group 5 septins like *Kp*AspE raise the possibility that this region may be used for more than just homomeric interactions. Intriguingly, the same R275E mutant that favors monomers over dimers also completely abrogates trimer formation at pH 6.4 (Figure 3). This suggests that perturbation of the G-interface disrupts both dimer and trimer formation. Canonical septin hetero-octamers form through a step-wise assembly pathway (Weems & McMurray, 2017); if a similar step-wise assembly pathway governs the formation of septin homo*-*oligomers, it is possible that dimer formation must precede trimer formation.

The family of septin proteins arose through repeated gene duplication and divergence, meaning that although all opisthokonts have multiple septin genes (Delic et al., 2024; Shuman & Momany, 2021) a single septin gene is the ancestral state of the septin cytoskeleton. As is the case with many heteromeric proteins (Mallik & Tawfik, 2020), heteromeric protein-protein interactions arose from ancestral homomeric interactions. Thus homomeric septin-septin interactions that persist in modern septins offer us a window into the evolutionary history of these cytoskeletal proteins. Such interactions include *Sc*Cdc10 dimers within septin octamers and *Sc*Cdc11 dimers between octamers. Of the canonical septins, Cdc11 is both most closely related to AspE (Figure 1) and the sole septin that interacts with it in yeast two-hybrid assays (Figure 5). But whereas Cdc10 and Cdc11 both form homodimers across NC-interfaces, AspE–– like the non-opisthokont septin *Cr*SEPT (Pinto et al., 2017)–– dimerizes across a G-interface that includes an arginine finger (Figures 1 & 3). The R-finger motif is widespread in non-opisthokont septins, present in only a small minority of Group 5 septins, and absent from septins in Groups 1-4, many of which have evolved into “pseudoGTPases” lacking the ability to either bind or hydrolyze GTP (Delic et al., 2024; Hussain et al., 2023). It is intriguing to note that Group 5 septins–– like non-opisthokont septins (Delic et al., 2024)–– appear to utilize G-interface residues for homodimer formation, while still retaining GTPase activity (Figure 2). But *Kp*AspE is also capable of interacting with other septins through an NC-interface that includes a ɑ0 helix, an ability and structural feature much more characteristic of opisthokont septins (Delic et al., 2024). Taken together, our findings suggest *Kp*AspE retains ancestral traits that are now characteristic of both opisthokont and non-opisthokont septins, consistent with the recent taxonomic assignment of Group 5 septins as the earliest-diverging clade of opisthokont septins (Delic et al., 2024).

It remains to be seen if filamentation is an ancestral septin trait. No filaments or higher-order structures were observed with recombinant *Kp*AspE *in vitro* (Supp. Figure 2). We conclude that either *Kp*AspE does not form filaments or additional factors–– perhaps proteins, small molecules, or post-translational modifications–– are necessary for their formation. In contrast to the lack of filamentation observed for *Kp*AspE, *Kp*Cdc12 definitively localizes to thick filaments *in vivo*, in both round and pseudohyphal cells (Figure 6). The *A. nidulans* homolog of Cdc12, AspC, also localizes as elongated filaments in hyphae (Lindsey et al., 2010). It remains to be seen if the other *K. petricola* septins will co-localize with *Kp*Cdc12, or if they will exhibit disparate localizations patterns, as was observed for the septin proteins of *A. fumigatus* (Juvvadi et al., 2013). Disruption of *A. fumigatus* microtubules with either nocodazole or benomyl eliminated formation of apparent *Af*AspE filaments (Juvvadi et al., 2013), suggesting that filamentation may not be an inherent property of Group 5 septins, but rather an apparent consequence of their interaction with other components of the cytoskeleton.

Because filamentation is not necessary for curvature sensing by septins (Cannon et al., 2019), we did not take the lack of filamentation *in vitro* to preclude curvature-sensing by *Kp*AspE. In an *in vitro* bead-binding assay, *Kp*AspE preferentially bound to the shallowest positive membrane curvature with which it was presented (Figure 4). This stands in contrast to canonical septin hetero-octamers, which prefer a somewhat higher micron-scale membrane curvature (Bridges et al., 2016). A C-terminal amphipathic helix in *Sc*Cdc12 is required for canonical septins to distinguish between different membrane curvatures (Cannon et al., 2019). HeliQuest predicts no similar amphipathic helix anywhere in *Kp*AspE (Gautier et al., 2008). Given that *K. petricola* cells grown on rich media are nearly spherical, presenting a uniform negative curvature to cytoplasmic proteins like septins, future studies should explore *Kp*AspE’s *in vitro* preference for negative membrane curvatures and *in vivo* localization patterns.

The biochemistry Group 5 septins such as *Kp*AspE has potential practical significance beyond deepening our understanding of the evolution of the cytoskeleton. A major challenge in the development of antifungal drugs is the relative similarity of animal and fungal cells. Septins have been implicated in the pathogenesis of multiple fungal pathogens of both plants and animals (Momany & Talbot, 2017). And drugs that disrupt septin assembly (by inhibiting the biosynthesis of very long chain fatty acids) have been successfully deployed to protect plants from fungal pathogens such as *Magnaporthe oryzae* (He et al., 2020). We argue that Group 5 septins, which are found exclusively in filamentous fungi (Delic et al., 2024), are of interest not only as windows into the evolutionary history of the septin cytoskeleton, but as potential targets for antifungal drugs.

The recombinant expression and biochemical characterization of a Group 5 septin lays the groundwork for future studies exploring the biology of this unique class of septins. Future work will explore interactions between Group 5 and canonical septins both *in vivo* and *in vitro*, as well as the regulation of *Kp*AspE by post-translational modifications and nucleotide binding state.

## Materials and Methods

### *Culture of* K. petricola

All *K. petricola* strains used in this study are derived from the sequenced A95 strain (CBS 123872). Unless otherwise stated, *K. petricola* cells were grown at 25°C on either rich media (YPD) or a synthetic complete media (ASM) that had been previously developed for *K. petricola* (Nai et al., 2013).

### Transformation of K. petricola

CRISPR-mediated transformation of *K.petricola* was performed as described by Voigt and colleagues (Voigt et al., 2020). Briefly, protoplasts were produced by overnight incubation of cells with a cocktail of lysing enzymes from *T. harzianum*, Yatalase^TM^ Enzyme (Takara Bio), and lyticase from *Arthrobacter luteus* (Sigma Aldrich). PEG-mediated transformation was used to introduce AMA1-containing vectors encoding guide RNAs and Cas9 (Nødvig et al., 2015; Wenderoth et al., 2017) and gene-replacement cassettes containing selectable markers (either hygromycin or nourseothricin).

### Rapid amplification of cDNA ends (RACE)

Total RNA was isolated from liquid cultures of *K. petricola* using the RNeasy Mini Kit (Qiagen, Catalog no. 74104). cDNA was synthesized and specific target amplified using the 3’ RACE System for Rapid Amplification of cDNA Ends (Invitrogen, Catalog no.18373-019) and 5’ RACE System for Rapid Amplification of cDNA Ends, Version 2.0 (Invitrogen, Catalog no. 18374-058). The gene-specific primers used are listed in Supplementary Table 1. The complete sequences of the *K. petricola* septin genes are listed in Supplementary Table 2.

### Recombinant expression and purification of KpAspE

#### Cloning

After RACE confirmed the locations of the N- and C-termini of the *KpAspE* gene, a codon-optimized synthetic gene product was synthesized in two overlapping halves by Twist Biosciences and integrated into a vector using the NEBuilder HiFi DNA Assembly Cloning Kit (NEB). pNH-TrxT was a gift from Opher Gileadi (Addgene plasmid # 26106) (Savitsky et al., 2010). This vector introduces cleavable N-terminal six-histidine and *E.coli* thioredoxin tags, for affinity purification and improved expression, respectively.

#### Expression

BL21 (DE3) *Escherichia coli* cells were transformed with the expression plasmid and selected for with kanamycin. Cells were grown in Terrific Broth (TB) at 37°C, 220 rpm, to an OD_600_ of 0.6–0.7. Expression was induced with 1 mM IPTG, and cells were grown at 18°C for 22 h. Cells were harvested by centrifugation at 13,689 x g for 15 min at 4°C. Pellets were resuspended in cold lysis buffer (50 mM HEPES, pH 7.4, 300 mM KCl, 1 mM MgCl_2_, 10% glycerol, 1% Tween 20 mM imidazole, 1 mM PMSF, 1 mg/mL lysozyme, and c0mplete^TM^ Protease Inhibitor Cocktail (Roche)) and either lysed immediately or stored at -80°C until lysis.

#### Cell lysis

Cells were thawed and sonicated on ice for a total of 10 min, in 1-min pulses separated by 2-min rests. The resulting whole-cell extract was clarified by centrifugation at 4°C for 30 min at 30,597 x g.

#### Ni^2+^ Affinity Purification

Clarified supernatant was loaded onto two sequential 5-mL HisTrap Crude FF columns (Cytiva), washed with 10 column volumes (CV) of Wash Buffer (50 mM HEPES, pH 7.4, 1 M KCl, 40 mM imidazole), and eluted in 25 mL of Elution Buffer (50 mM HEPES, pH 7.4, 300 mM KCl, 500 mM imidazole).

#### Size exclusion chromatography (SEC)

The HisTrap column elution was concentrated to a final volume of 1 mL using 30-kDa MW cutoff Amicon Ultra centrifugal filters (Millipore). The entire 1 mL was injected onto a Superdex 200 Increase 10/300 GL column (Cytiva) equilibrated SEC Buffer (50 mM HEPES, pH 7.4, 300 mM KCl, 1 mM BME). 6xHis-TxrT-*Kp*AspE typically eluted at about 12 mL, consistent with the predicted molecular weight of a dimer (146 kDa) (Figure 1).

#### Tag removal, dialysis, and storage

The SEC peak elution fractions were pooled in a 20-kDa MW cutoff cassettes (Thermo Fisher Scientific), incubated with ProTEV Plus protease (Promega), and dialyzed into Septin Storage Buffer (50 mM HEPES, pH 7.4, 300 mM KCl, 1 mM BME, 5% glycerol) overnight at 4°C via two 500-ml steps.

Protein was then run over 2-mL Ni^2+^-NTA resin bed to remove cleaved 6xHIS-TrxT tag, and TEV protease. Purity was assessed by SDS-PAGE using pre-cast Any kD^TM^ Mini-PROTEAN® TGXTM polyacrylamide gels (Biorad). Aliquots were flash-frozen and immediately stored at - 80°C.

Protein concentration was determined by Bradford or bicinchoninic acid (BCA) assay.

#### Atto NHS Ester Labeling

Atto 488 (Sigma Aldrich) was incubated with purified KpAspE at a 3-fold excess molar ratio and nutated at room temperature for 45-60 min. Excess dye was removed using Zeba^TM^ 40-kDa MW cutoff desalting columns (Thermo Scientific).

### Mass photometry

A ReFeyn One^MP^ instrument was used to measure the distribution of molecular masses of purified *Kp*AspE protein at 1-10 nM concentrations in SEC Buffer of varying pH. Refeyn DiscoverMP software was used for preliminary data analysis.

### Nucleotide binding assay

Purified *Kp*AspE protein was combined with a BODIPY-labeled guanosine nucleotide (Thermo Fischer). A CLARIOstar^PLUS^ plate reader was used to monitor fluorescence emission at 515 nm.

### Nucleotide hydrolysis assay

Purified KpAspE incubated with a 10-fold molar excess of guanosine nucelotide. The mixture was nutated at room temperature. Hydrolysis reactions were halted by flash-freezing in liquid N_2_, followed by boiling at 100°C for 10 minutes to denature all proteins and centrifugation to pellet the denatured protein. The relative abundance of different guanosine nucleotide species in each supernatant was analyzed using reverse-phase high-pressure liquid chromatography (HPLC) on a Hypersil™ ODS C18 column (Thermo Fischer; Catalog number 30105-052130) under isocratic conditions in running buffer (10 mM tetrabutyl ammonium bromide (TBAB), 100 mM Na_3_PO4, 5% acetonitrile, pH 6.5).

### Curvature sensing assay

*Kp*AspE protein adsorption onto silica microspheres of varying diameters coated in supported lipid bilayers was measured as previously described (Cannon et al., 2019; Woods et al., 2021).

### Yeast two-hybrid assay

Yeast two-hybrid assays were performed as previously described (Fields & Song, 1989). Briefly, the *Saccharomyces cerevisiae* reporter strain PJ69-4α was co-transformed with a plasmid containing GAL4 activation domain fused to a *Knufia petricola* septin (LEU2 selection) and a second plasmid containing GAL4 DNA-binding domain fused to a second *K. petricola* septin (TRP selection). In order to test the GAL4-driven transcription of the HIS3 reporter, transformants were plated on synthetic complete media minus histidine and containing various concentrations of 3-amino-1,2,4-triazole (3-AT), a competitive inhibitor of the HIS3 gene product.

### Cell microscopy

*K. petricola* cells were grown in liquid ASM at 25°C and either fixed using 3.7% formaldehyde or imaged live. For staining with calcofluor white, cells were incubated at room temperature with 0.01% calcofluor white in 1X PBS for 10 minutes and then washed twice with 1X PBS before mounting and imaging. Cells were imaged at 405 nm on a spinning disc confocal microscope Nikon CSU-W1 with a VC Plan Apo 100X/1.49 NA oil (Cargille Lab 16241) immersion objective and an sCMOS 85% QE 95B camera (Photometrics).

Whole colonies were grown on solid media at 25°C. Chunks of agar containing colonies were excised from the petri dish using a scalpel, then inverted in chambered coverslips (Ibidi) containing 50 µL of synthetic complete media. Strains containing GFP-labeled proteins were imaged at 510 nm on a confocal microscope Zeiss LSM 980 with Airyscan 2.

### AlphaFold Predictions

AlphaFold predictions were executed using the Colabfold Google notebook v1.5.5. Predictions primarily used an MMseqs2 MSA. Five models with three recycles each were generated and the highest ranking model was selected. The resulting 3D structures were visualized using ChimeraX(Meng et al., 2023) and colored according to protein identity.

### Data analysis and visualization

For analysis of septin binding, raw images were exported into Imaris 8.1.2 (Bitplane AG). All other data were analyzed in R (R Core Team, 2013). Figures were produced using Adobe Illustrator and the ggplot2 package in R (Wickham, 2009).

## Abbreviations

List only nonstandard abbreviations that are used three or more times in the text.

## Acknowledgements

We thank Leiah Marie Carey (Campbell Lab, UNC) for her instruction and support in the use of HPLC. We thank Nathan Nicely (Protein Expression and Purification Core Facility, UNC) for instruction and support in the use of mass photometry, as well as the use of a FPLC.

We thank Ellysa Vogt and Brandy Curtis for instruction and support in biophysical assays of septin behavior.

We are grateful to the entire Gorbushina Lab (BAM), especially Julia Schumacher, for sharing their *Knufia petricola* genetic toolkit, including a draft genome of the A95 lab strain.

G.E.H is funded by a NIGMS IRACDA fellowship through the SPIRE program at UNC. This work is supported by NSF Grant 2016022 to A.S.G.

---

**Supplementary figure 1.**
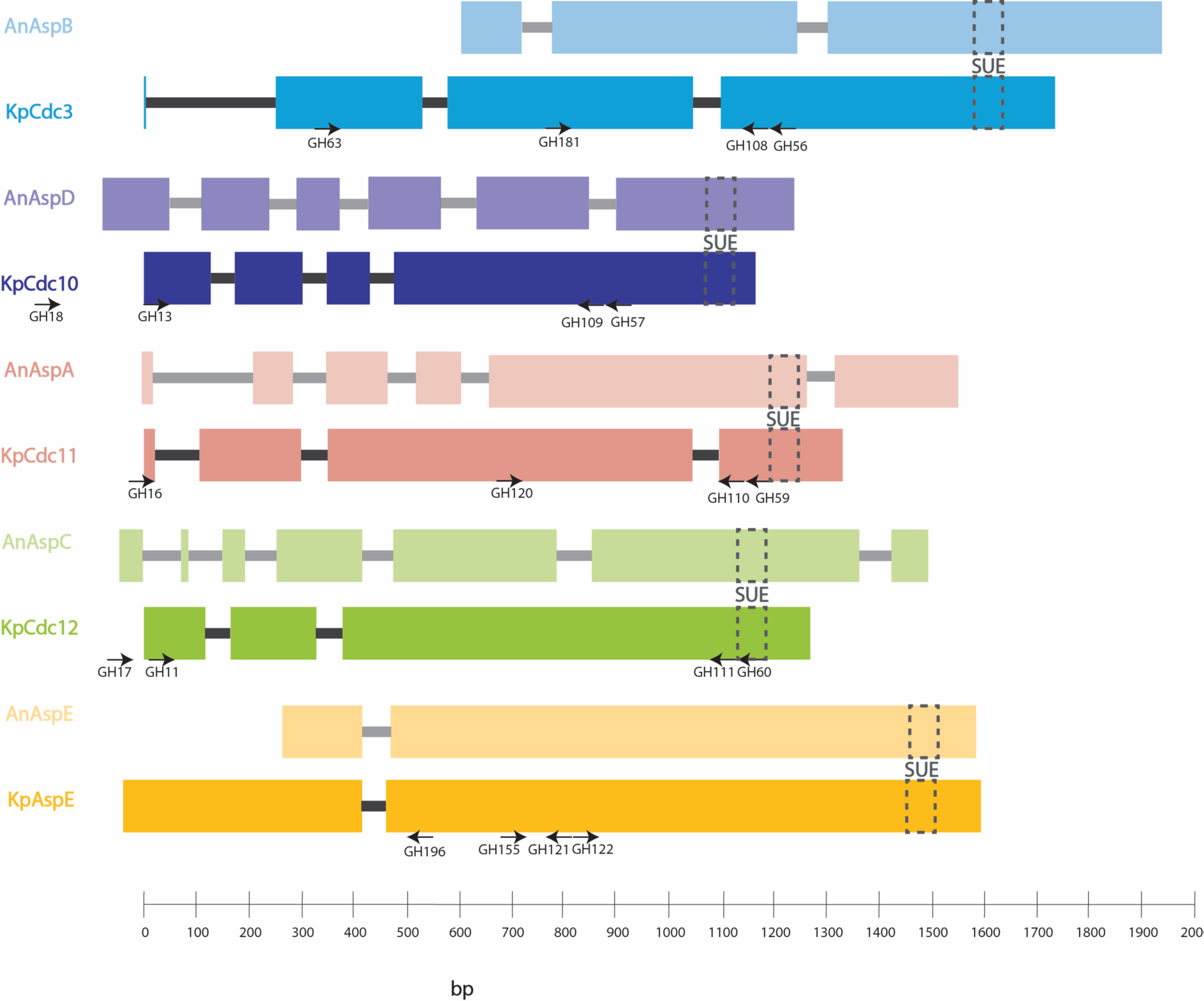
Structure and products of the five *Knufia petricola* septin genes. Intron-exon structure of the five *K. petricola* and *A. nidulans* septin genes. The locations of gene-specific primers used for rapid amplification of cDNA ends are indicated by black arrows.

**Supplementary table 1.**
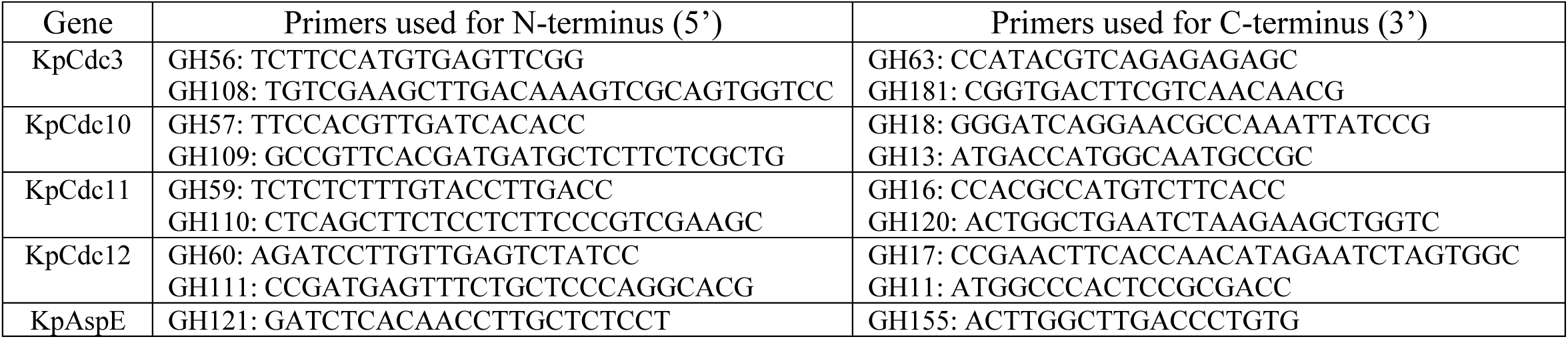
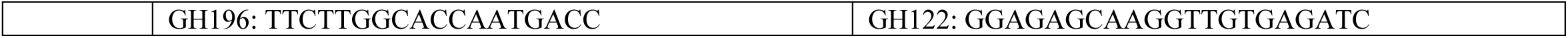
Gene-specific primers used for rapid amplification of cDNA ends (RACE).

**Supplementary table 2.**
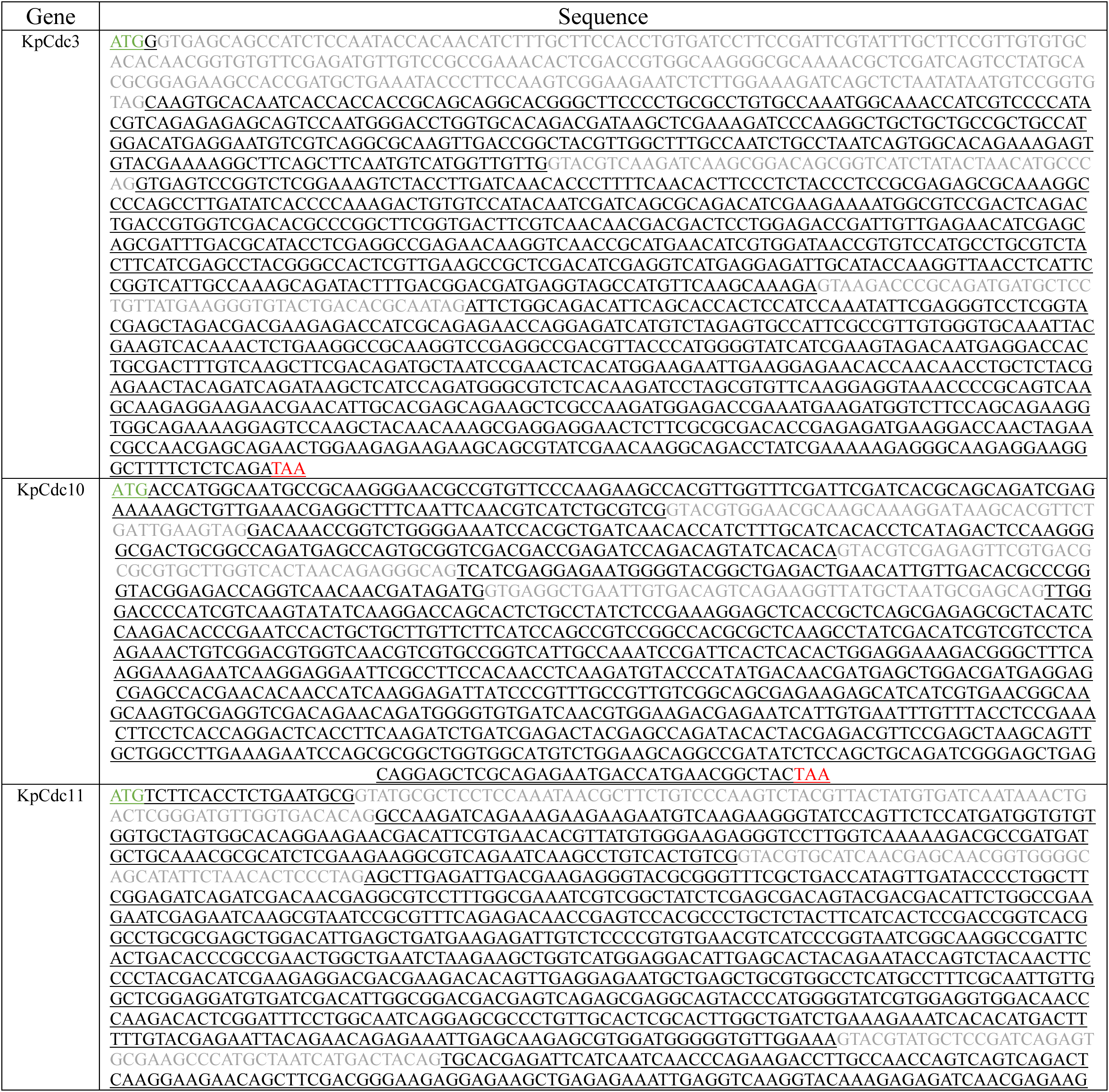
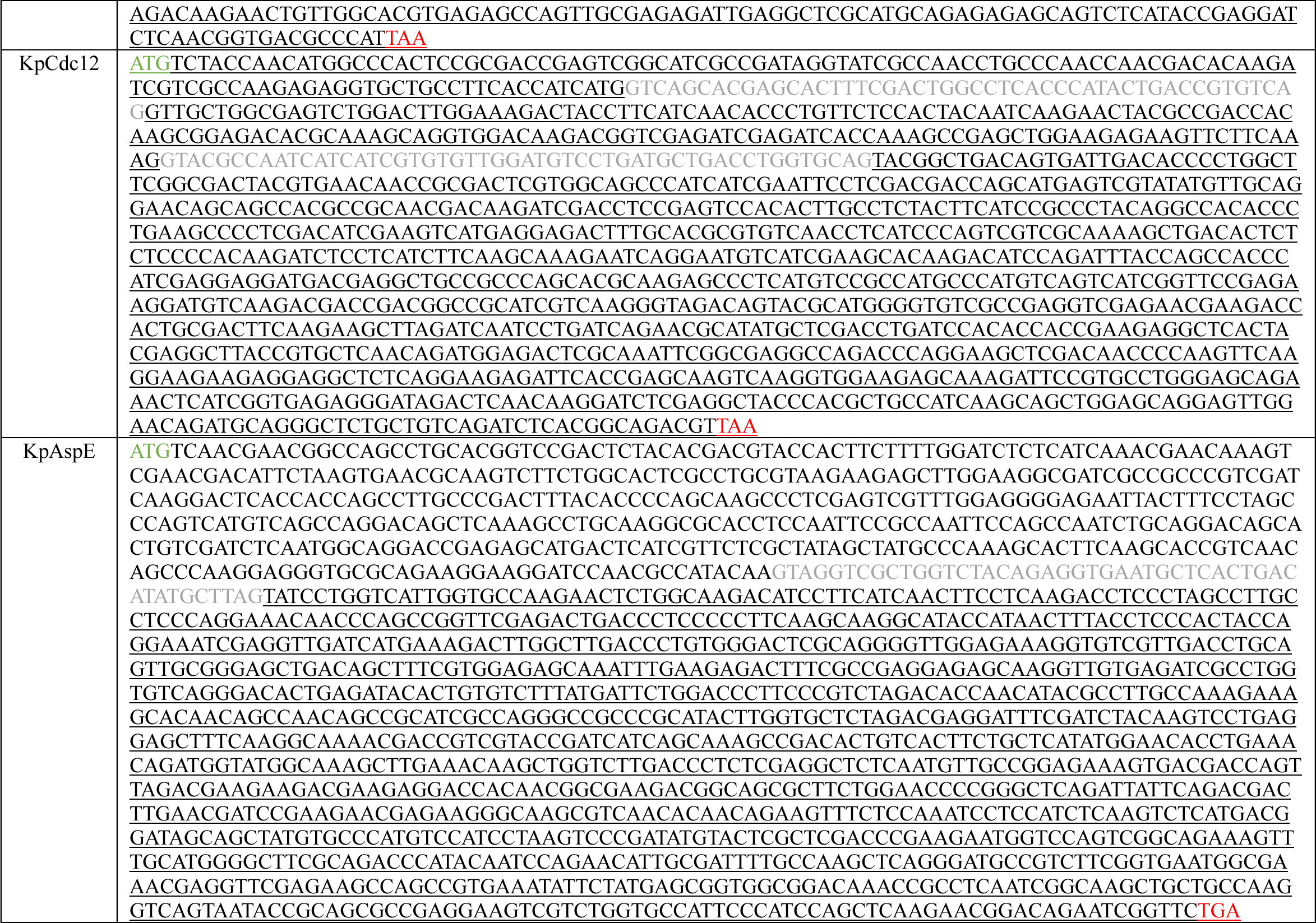
Sequences of *K. petricola* septin genes. Introns are shown in gray; exons are shown in black and underlined. Start codons are shown in green; stop codons are shown in red.

**Supplementary figure 2.**
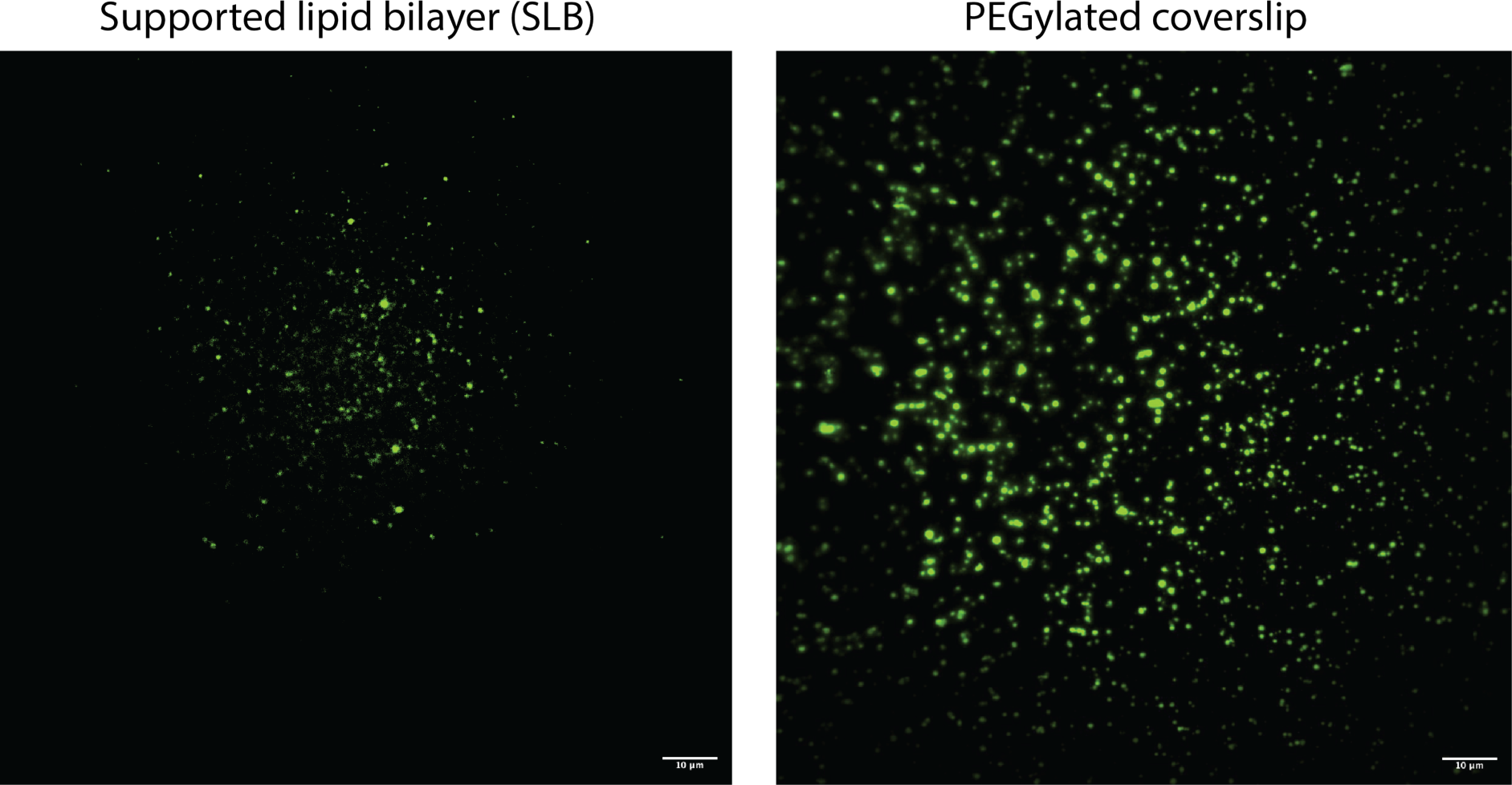
*Knufia petricola* AspE homo-oligomers do not spontaneously assemble into filaments under conditions that cause filamentation of *S. cerevisiae* canonical septin octamers. Purified, Atto488-labeled *Kp*AspE was incubated on either (A) supported lipid bilayers or (B) PEGylated coverslips at concentrations ranging from 5 to 100 nm, as previously described (Khan et al., 2018). No filamentation was ever observed on either substrate.

**Supplementary figure 3.**
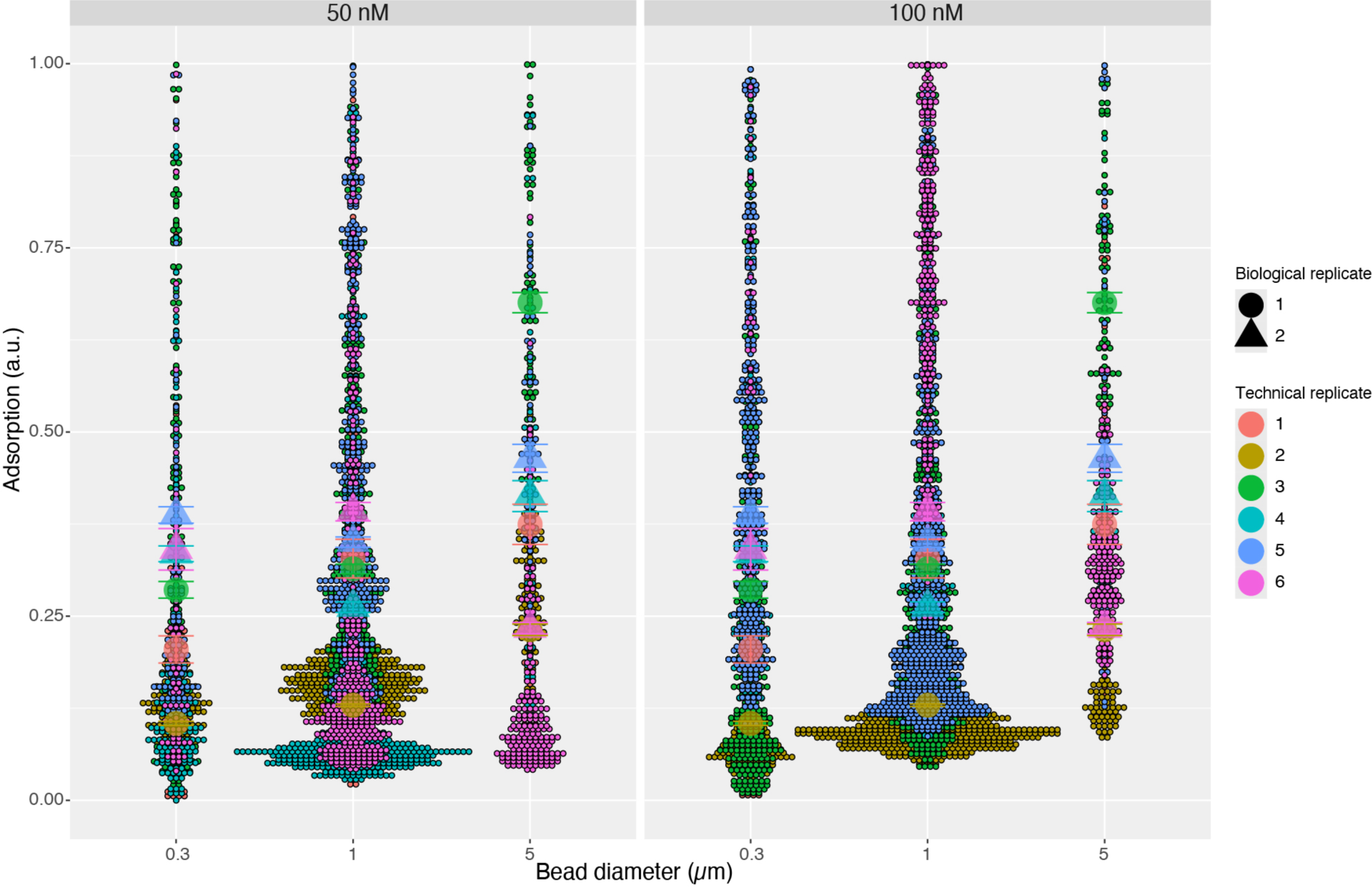
*in vitro Kp*AspE preferentially localizes to shallow positively curved membranes. Superplot of *Kp*AspE adsorption levels observed for beads of different sizes. Technical replicates are indicated by dot color. Large shapes indicate means for each technical replicate, while error bars extend to mean ± standard error. We consider recombinant proteins expressed and purified from separate populations of bacteria as biological replicates, which are indicated by dot shape. In five out of six technical replicates, *Kp*AspE adsorption was greatest on the beads with the largest diameter. We attribute the anomalous result in technical replicate six to variability in the ratio of beads of different sizes; on that day, 5-μm beads were 34% of the total bead population, much higher than their average abundance of 15%. Previous work has demonstrated that the curvature preference of septins is sensitive to the combination of membrane curvatures present in a reaction (Shi et al., 2023). Thus, variability in the ratio of bead sizes is a source of technical variation in this assay.

**Supplementary figure 4.**
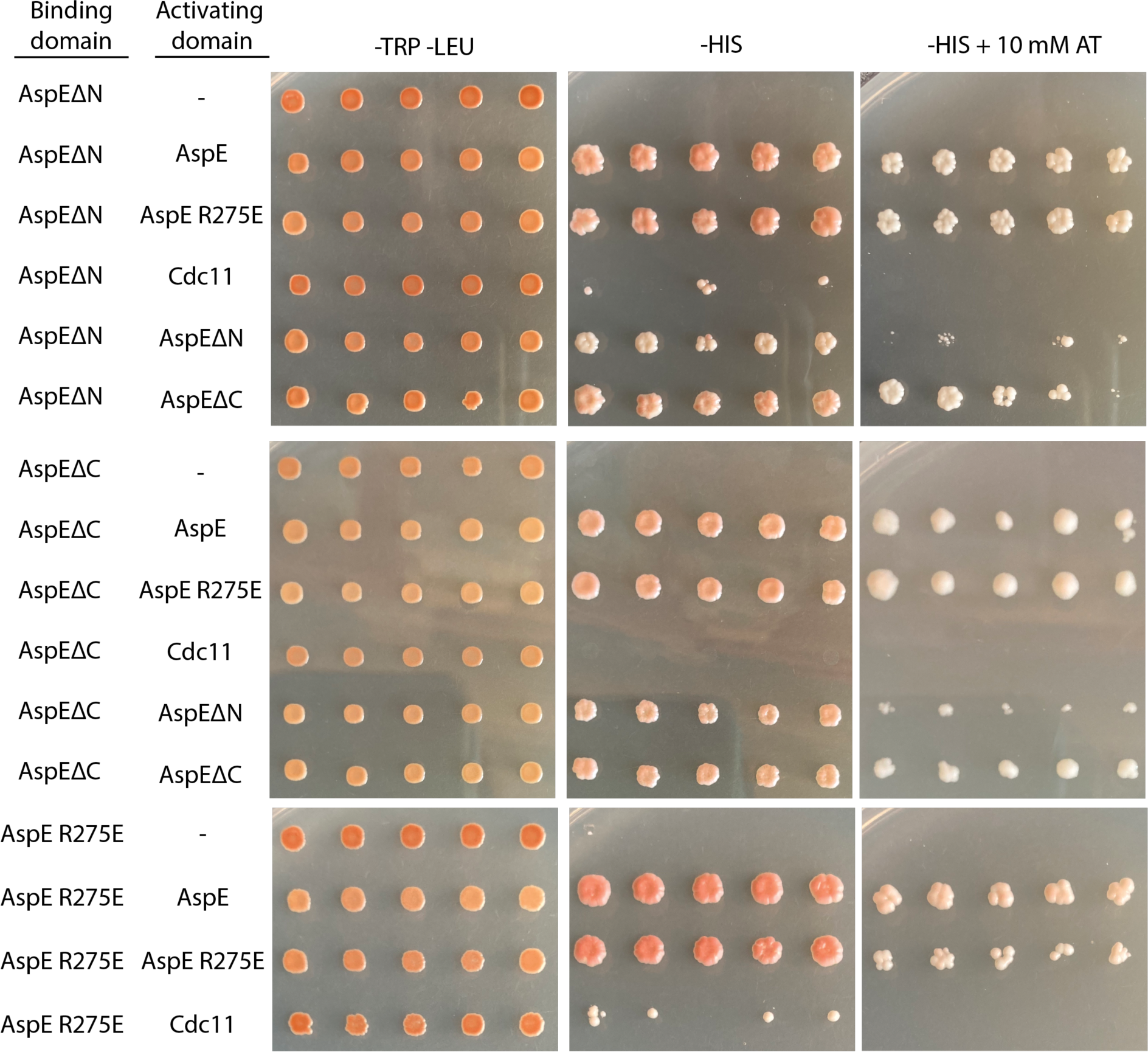
The N- and C-termini of *Kp*AspE are necessary for its interaction with *Kp*Cdc11 but not for interaction with itself.

